# Structural genomic applied on the rust fungus *Melampsora larici-populina* reveals two candidate effector proteins adopting cystine-knot and nuclear transport factor 2-like protein folds

**DOI:** 10.1101/727933

**Authors:** Karine de Guillen, Cécile Lorrain, Pascale Tsan, Philippe Barthe, Benjamin Petre, Natalya Saveleva, Nicolas Rouhier, Sébastien Duplessis, André Padilla, Arnaud Hecker

## Abstract

Rust fungi are plant pathogens that secrete an arsenal of effector proteins interfering with plant functions and promoting parasitic infection. Effectors are often species-specific, evolve rapidly, and display low sequence similarities with known proteins or domains. How rust fungal effectors function in host cells remains elusive, and biochemical and structural approaches have been scarcely used to tackle this question. In this study, we used a strategy based on recombinant protein production in *Escherichia coli* to study eleven candidate effectors of the leaf rust fungus *Melampsora larici-populina.* We successfully purified and solved the three-dimensional structure of two proteins, MLP124266 and MLP124017, using NMR spectroscopy. Although both proteins show no sequence similarity with known proteins, they exhibit structural similarities to knottin and nuclear transport factor 2-like proteins, respectively. Altogether, our findings show that sequence-unrelated effectors can adopt folds similar to known proteins, and encourage the use of biochemical and structural approaches to functionally characterize rust effector candidates.

## INTRODUCTION

To infect their host, filamentous pathogens secrete effector proteins that interfere with plant physiology and immunity to promote parasitic growth (Win *et al.*, 2012). Although progresses have been made in the past decade, how effectors act in host cells remains a central question in molecular plant pathology. Effectors of filamentous pathogens are secreted and either stay in the apoplast or penetrate inside the cell cavity through specialized infection structures such as haustoria (Petre and Kamoun, 2014). Effectors are detected by the host plant by two layers of immune receptors at the cell surface or inside the cell, which triggers plant defence response (Jones and Dangl, 2006).

To evade recognition by the host immune system, pathogen effector genes evolve rapidly, notably through the diversification of the amino acid sequence of the encoded proteins (Persoons *et al.*, 2017). Such diversification impairs the identification of amino acid motifs or sequences similar to known proteins, which could give insights on effectors function inside the host cell (Franceschetti *et al.*, 2017). However, effector protein structures may be more conserved than their primary sequences. Indeed, several superfamilies of effector proteins, such as the fungal MAX or the oomycete WY-domain families, have members showing similar fold but divergent primary sequences (de Guillen et al. 2015; Win et al. 2012) the fold being conserved probably due to the strong link between protein structure and function (Illergård *et al.*, 2009). Thus, structure resolution of candidate effectors represents a powerful approach to gain insights into effector functions, but also to define and identify structural effector families. Research efforts have been recently set in this direction and applied in order to determine the structure of several effector proteins (de Guillen *et al.*, 2015; Ve *et al.*, 2013; Wang *et al.*, 2007; Wirthmueller *et al.*, 2013).

Rust fungi (Pucciniales, Basidiomycetes) constitute the largest group of obligate biotrophic pathogens, and they infect almost all plant families, causing serious damages to cultures (Dean *et al.*, 2012; Lorrain *et al.*, 2019). In the past decade, progresses made in rust fungal effector biology were mostly based on genomics and transcriptomics, and common but not exclusive features have been used to identify candidate effectors in predicted secretomes (Duplessis *et al.*, 2014). Catalogues of hundreds to thousands of candidate effectors have been unravelled. Generally, criteria such as the protein size, the presence of a predicted secretion signal, the absence of functional information, the richness in cysteines, the transcriptional regulation during infection, and/or the presence of signatures of rapid evolution, have been used as a basis for identifying candidate effectors (Cantu *et al.*, 2013; Hacquard *et al.*, 2012; Nemri *et al.*, 2014; Saunders *et al.*, 2012). Due to the difficulty to genetically manipulate rust fungi and their host plants, only a handful of rust effectors have been characterized so far (Petre and Kamoun, 2014). Apart from their avirulence properties (i.e. recognition by plant immune receptors inside the cell), the functions of these rust effectors remain unknown (Lorrain *et al.*, 2019). Effectoromic pipelines have been recently established to prioritize rust candidate effectors based on heterologous systems to get insights about their plant cellular and molecular targets (Germain *et al.*, 2018; Lorrain *et al.*, 2018; Petre *et al.*, 2015; Petre *et al.*, 2016; Qi *et al.*, 2018). But so far, only one study has set up a small-scale effort using production of candidate effectors in bacterial system to unravel their structure and function (Zhang *et al.*, 2017). The identification of plant targets of effectors associated with structure/function analyses of recombinant effectors can reveal how they interact with plant partners and how co-evolution with the plants promotes the diversification of surface-exposed amino acids (Boutemy *et al.*, 2011; Chou *et al.*, 2011; Leonelli *et al.*, 2011; Wang *et al.*, 2007; Win *et al.*, 2012; Yaeno *et al.*, 2011). The avirulence proteins AvrL567, AvrM, and AvrP of the flax rust fungus *Melampsora lini* are the three effector structures described in rust fungi so far (Ve *et al.*, 2013; Wang *et al.*, 2007; Zhang *et al.*, 2018).

The poplar rust fungus *Melampsora larici-populina* is the causal agent of the poplar leaf rust disease. It causes important damages in poplar plantations across Europe (Pinon and Frey, 2005). It is also a model pathosystem to study tree-microbe interaction. As such, recent research efforts have identified and initiated the characterization of *M. larici-populina* candidate effectors using transcriptomics and functional screens in heterologous plant systems such as *Nicotiana benthamiana* and *Arabidopsis thaliana* (Gaouar *et al.*, 2016; Germain *et al.*, 2018; Petre *et al.*, 2015; Petre *et al.*, 2016). These studies highlighted that *M. larici-populina* candidate effectors target multiple cell compartments and plant proteins. The same conclusions have been drawn from similar effectoromic screens in other rust fungi (Lorrain *et al.*, 2018). It remains to make progress in the determination of the functions of unravelled priority effectors.

In this study, we combined biochemical and structural approaches to explore further *M. larici-populina* candidate effector proteins. To this end, we used *Escherichia coli* as an heterologous system to express eleven candidate effectors that were previously described to target particular cell compartments and/or to interact with specific plant proteins and/or that are homologues of known rust avirulence effectors (Gaouar *et al.*, 2016; Germain *et al.*, 2018; Petre *et al.*, 2015; Petre *et al.*, 2016). Among the eleven selected proteins, only three were successfully produced and purified from *E. coli* as recombinant proteins. We could determine the nuclear magnetic resonance (NMR) structures of two of them, highlighting structural similarities with Nuclear Transport Factor 2-like proteins and with Knottins.

## RESULTS

### Selection of *M. larici-populina* candidate effector proteins

For this study, we selected eleven candidate effector proteins of *M. larici-populina* (Table 1). These proteins are a subset of a previous catalogue of >25 candidate effectors that have been screened *in planta* for their subcellular localization and plant protein partners (Lorrain *et al.*, 2018; Petre *et al.*, 2015). These selected candidates are all small-secreted proteins that are expressed *in planta* or in haustoria, specific to rust fungi, with no known function, or with homologies to known rust effector proteins. Notably, we retained proteins showing i) a specific and informative subcellular localization, such as nucleus (MLP109567), nucleolus (MLP124478), nuclear bodies (MLP124530), chloroplasts and mitochondria (MLP107772, aka CTP1), chloroplasts and aggregates (MLP124111), endomembranes (MLP124202), and plamodesmata (MLP37347), ii) specific plant partners (MLP124017, MLP37347, MLP124448, MLP124111), iii) similarities with *M. lini* Avr effectors (MLP124530, MLP37347, MLP124202, MLP124266), or iv) belonging to large families of small-secreted proteins (MLP124499, MLP124561).

**Table 1:**
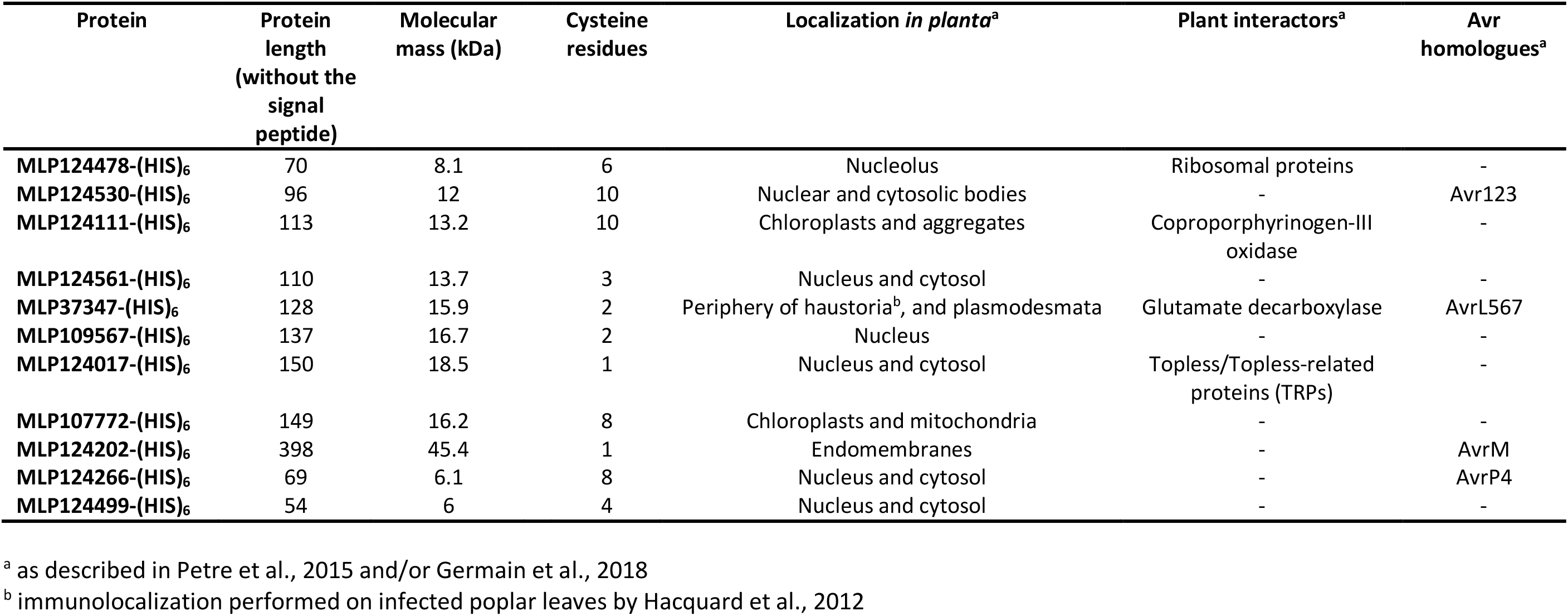
Features of the eleven *M. larici-populina* candidate effector proteins investigated in this study

### Three out of eleven candidate effectors were successfully produced in *E. coli* and purified

To investigate the structural properties of the 11 selected candidate effectors (Figure 1), we first aimed at obtaining the recombinant proteins. To this end, we first cloned the sequence coding the mature proteins (i.e. without signal peptide) into pET-26b (for *Mlp124111*, *Mlp124478*, *Mlp124530*, *Mlp124561*, *Mlp37347*, *Mlp109567*, *Mlp107772*, and *Mlp124202*) or pET-28a (for *Mlp124266*, *Mlp124499*, and *Mlp124017*) expression vectors, in order to incorporate a C-terminal 6-histidine tag that allows protein purification by immobilized metal ion affinity chromatography (IMAC) (Table S1). We then inserted the expression vectors into the *E. coli* Rosetta2 (DE3) pLysS strain and induced the production of the His-tagged recombinant proteins. Small-scale expression assays indicated that nine out of the eleven proteins accumulated using a standard induction protocol (i.e. addition of 100 µM IPTG in mid-exponential growth phase and further growing for 3 to 4 hours at 37°C). Among those, five (MLP124111, MLP124561, MLP37347, MLP107772 and MLP124202) accumulated in the insoluble protein fraction, five (MLP124478, MLP124530, MLP124017, MLP124266 and MLP124499) were expressed as soluble proteins and one (MLP109567) is not expressed (Figure 2). Despite the use of other *E. coli* expression strains (SoluBL21 (DE3), Origami2 (DE3) pLysS, as well as Rosetta-Gami2 (DE3) pLysS) which are known to increase the solubility of recombinant proteins or to improve the maturation of disulfide bridges, and despite the modification of the induction conditions (induction time, temperature, and osmolarity), we were neither able to enhance the expression of MLP109567 nor the solubility of MLP124111, MLP124561, MLP37347, MLP107772 and MLP124202. As MLP124017, MLP124266, and MLP124499 were expressed in the soluble protein fraction of *E. coli* and enough stable along the purification procedure, we decided to retain these three proteins for further analyses. We thus purified the His-tagged recombinant proteins in native conditions using a two-step protocol including first affinity chromatography, then exclusion size chromatography (Figure 3). The purified proteins, yielding respectively 50 mg/L (cell culture), 0.5 mg/L and 0.5 mg/L for MLP124017, MLP124266 and MLP124499, respectively, eluted in size exclusion chromatography as a single peak corresponding to an estimated apparent molecular mass compatible with a monomeric organization.

**Figure 1:**
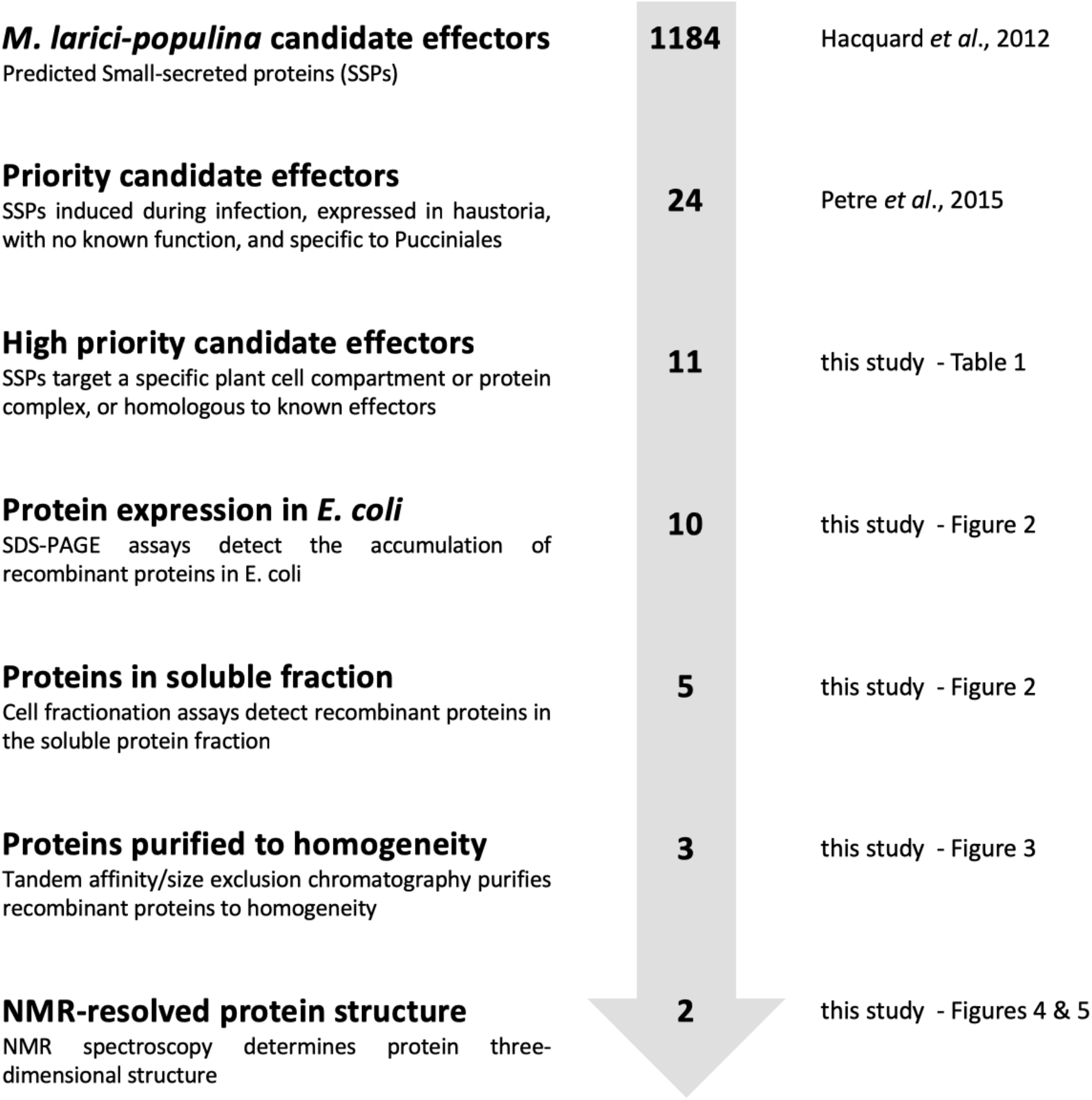
Overview of the effectoromics pipeline. A total of eleven *M. larici-populina* candidate effectors selected from the previous study of Petre et al. (2015) (i.e. particular localization and/or specific plant interactors and/or homologies to *M. lini* Avr effectors). Effector candidates were expressed in *E. coli* SoluBL21 (DE3) pRARE2, Rosetta2 (DE3) pLysS or RosettaGami2 (DE3) pLysS strains. Soluble recombinant proteins were purified and their structure solved by NMR spectroscopy.

**Figure 2:**
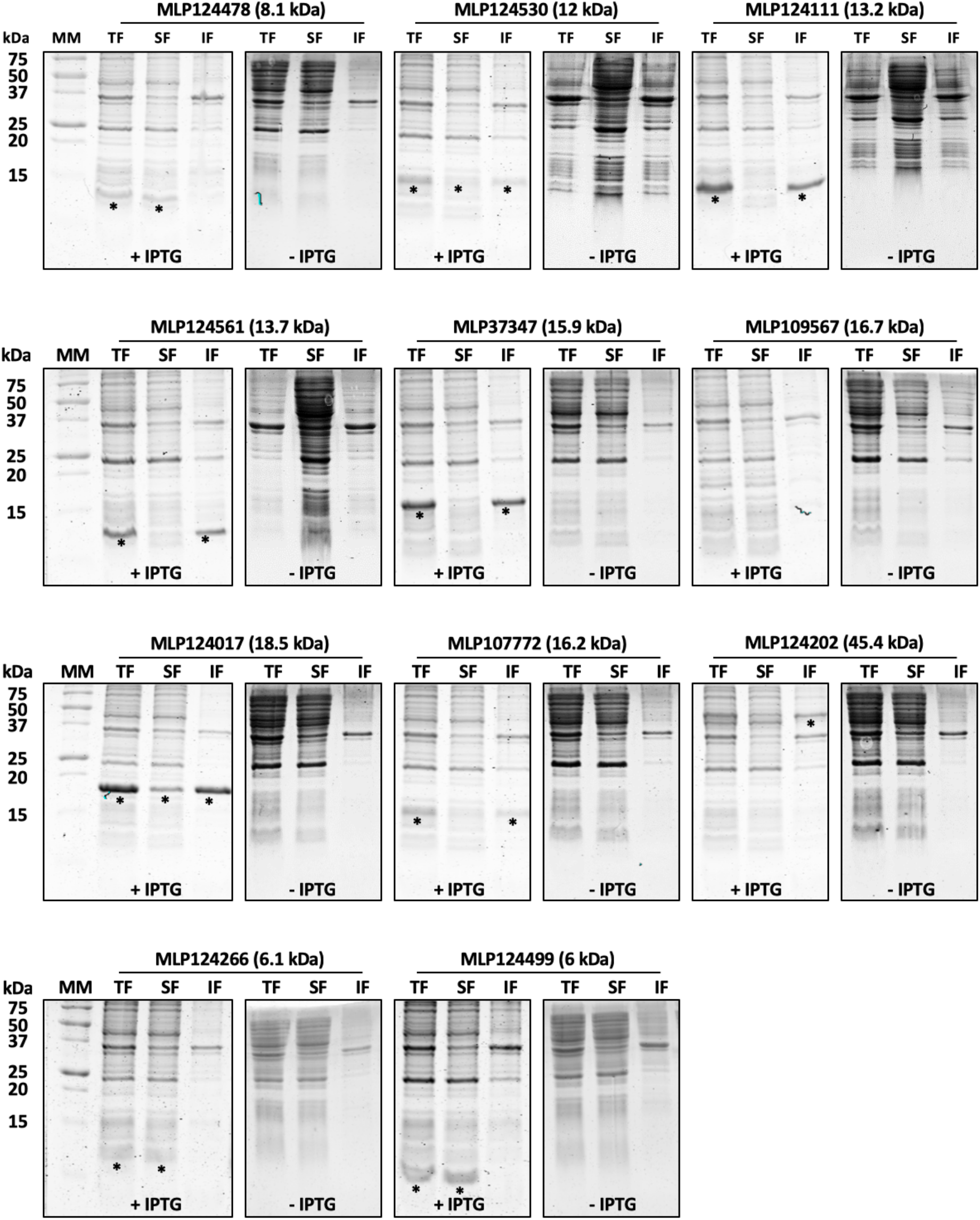
Small-scale expression test of selected candidate effector proteins carried out in *Escherichia coli* Rosetta2 (DE3) pLysS expression strain. Coomassie blue-stained sodium dodecylsulfate-polyacrylamide gel electrophoresis (SDS-PAGE) analysis of total (TF), soluble (SF) and (IF) insoluble protein fractions of *E. coli* Rosetta2 pLysS expression strain grown in presence (+) or in absence (-) of 0.1 mM isopropyl β-D-thiogalactopyranoside (IPTG). Asterisks indicate the expected migration of overexpressed proteins. MM: molecular mass marker.

**Figure 3:**
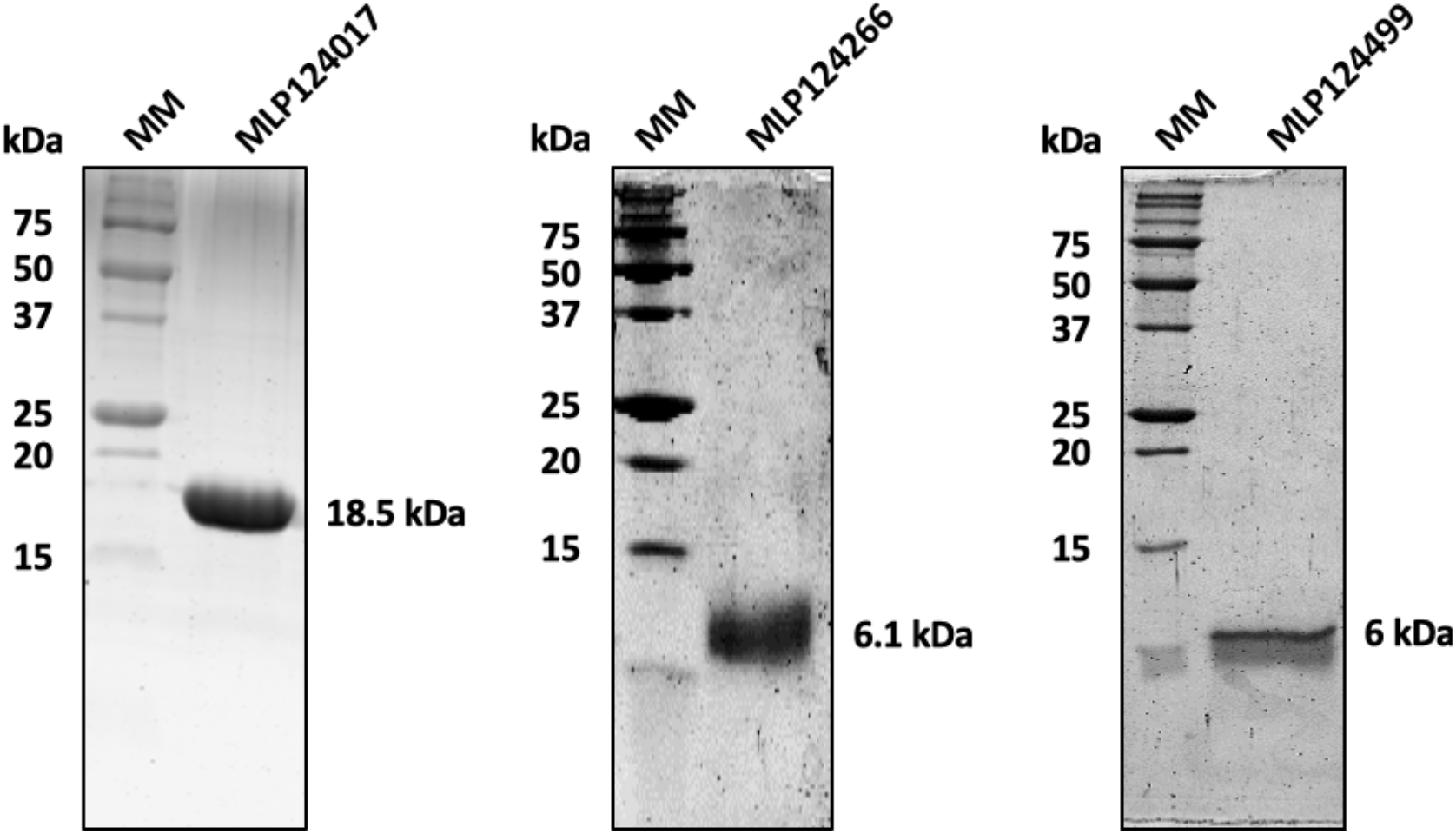
Candidate effectors purified as soluble proteins. Ten micrograms of recombinant MLP124017-(His)_6_; MLP124266-(His)_6_, MLP124499-(His)_6_ have been separated by SDS-PAGE (17%). Molecular mass corresponding to each purified proteins is indicated. MM: molecular mass marker.

### MLP124266 is a thermosoluble protein that exhibits a cystine-knot

From a previous study, we reported that the Mlp124266 and Mlp124499 genes are strongly expressed and induced during poplar leaf colonization by *M. larici-populina*, and belong to large multigene families of 13 and 31 members, respectively, in *M. larici-populina* (Hacquard *et al.*, 2012). Mlp124266 and Mlp124499 encode mature proteins of 69 and 50 amino acids, respectively (Figure S1A). MLP124266 has an N-terminal part enriched in charged residues and a C-terminal region that possesses six conserved cysteines predicted to form a cystine-knot structure. This typical protein organization is shared by all members of the family as well as by alleles of *M. lini* AvrP4 (Barrett *et al.*, 2009; Persoons *et al.*, 2014). In MLP124499, several acid residues are found in the N-terminal part whereas the C-terminal part contains three conserved cysteines (Figure S1B). Prediction programs indicate that all members of both protein families exhibit highly conserved N-terminal signal peptides for protein secretion. Following the production and the purification of both MLP124266 and MLP124499, we undertook a structural characterization of each recombinant protein using a NMR spectroscopy approach.

Standard homonuclear 2D experiments and ^15^N-edited TOCSY-HSQC and NOESY-HSQC experiments carried out on MLP124499 (Table S2) allowed the assignment of ^1^H and ^15^N resonances except for the four N-terminal residues (Figure S2B). However, several minor peaks were observed, especially for Ala_14_-Glu_16_, Gly_25_-Gln_26_, Glu_30_, Asp_49_ residues, suggesting the presence of multiple forms or conformations (Figure S2A). Changing the temperature and the ionic strength, or adding dithiothreitol failed to improve the quality of the MLP124499 NMR spectra. Therefore, the structural models obtained were not well enough defined to characterize a specific fold for this protein.

For MLP124266, the assignment of ^1^H, ^15^N and ^13^Cresonances has been obtained for all residues except the five N-terminal amino acids (Figure S2A) and its 3D structure could further be modelled by the NMR derived constraints (Figure 4; Figure S3). This analysis showed that the Cys_36_-Leu_69_ C-terminal region exhibits a typical cystine-knot structure involving three disulfide bonds (Cys_39_-Cys_55_, Cys_44_-Cys_58_, and Cys_50_-Cys_64_), a β-sheet composed of anti-parallel strands between Thr_42_-Cys_44_, Gly_57_-Ser_59_ and Val_63_-Val_65_, and a short α-helix formed by the Gln_49_-Ala_52_ segment. In contrast, the Met_1_-Asp_35_ N-terminal region displays large structural disorder, as shown by the superposition of the 20 NMR models (Figure S3). Very few NOE correlations were indeed observed for residues 1 to 35. A few sequential and medium-range NOE correlations characteristic of transient helical conformations can however be noticed (Figure S3) and explain the presence of short secondary structures in some of these 20 models. Indeed, residues 8-17 and 28-30 exhibit helical structures in 30 to 50 % and around 25 % of the models, respectively. Moreover, backbone dynamic properties of MLP124266 have been investigated by ^15^N relaxation measurements. Heteronuclear ^1^H-^15^N NOE values showed a contrasted profile with low values for N-terminal residues (indicative of a flexible structure) and high values for C-terminal residues (indicative of a rigid structure). Indeed, amino acids Asp_6_ to Gly_38_ and Cys_39_ to Leu_69_ presented heteronuclear NOE averaged values of 0.26 ± 0.05 and 0.66 ± 0.10, respectively. Interestingly 6 out of the 8 cysteines are gathered in the C-terminal region between Cys_39_ and Cys_69_, following a spacing (Cys-X_2-7_-Cys-X_3-10_-Cys-X_0-7_-Cys-X_1-17_-Cys-X_4-19_-Cys) typical of cystine-knot structures, (i.e., three intricate disulfide bridges that confer very high stability to proteins; Figure 4; (Daly *et al.*, 2011)). Hence, it is likely that rigidity originates from the structure formed by these cysteines that are highly conserved in the protein family, as indicated by a Consurf analysis (Figure S1, Figure S4). Thus, we sought to determine whether these disulfides are formed and whether they influence the stability and/or the oligomerization state of the protein by covalent bonds. A single peak corresponding to the theoretical mass of MLP124266 monomer was obtained by mass spectrometry (data not shown). The titration of free thiol groups in an untreated recombinant MLP124266 gave an averaged value of 1 mole SH per mole of protein. Considering the presence of 8 cysteines in the protein, these results are consistent with the existence of three intramolecular disulfide bridges (Figure S3). The thermostability of recombinant protein was estimated by heating the protein for 10 min at 95°C. The fact that the protein remained in solution (*i.e.* no precipitation was observed) led us to conclude that it is thermosoluble. In order to investigate the role of the disulfides for such property, we should compare the results obtained with an oxidized and a reduced protein. However, as assessed by thiol titration experiments, we failed to obtain a complete reduction of these disulfides despite extensive incubation of the protein at high temperatures, in denaturing and reducing environments. Altogether, these results indicate that recombinant MLP124266 is properly folded by *E. coli*, and that the disulfide bridges, which are somehow resistant to reduction, confer a high rigidity and stability to the protein.

**Figure 4:**
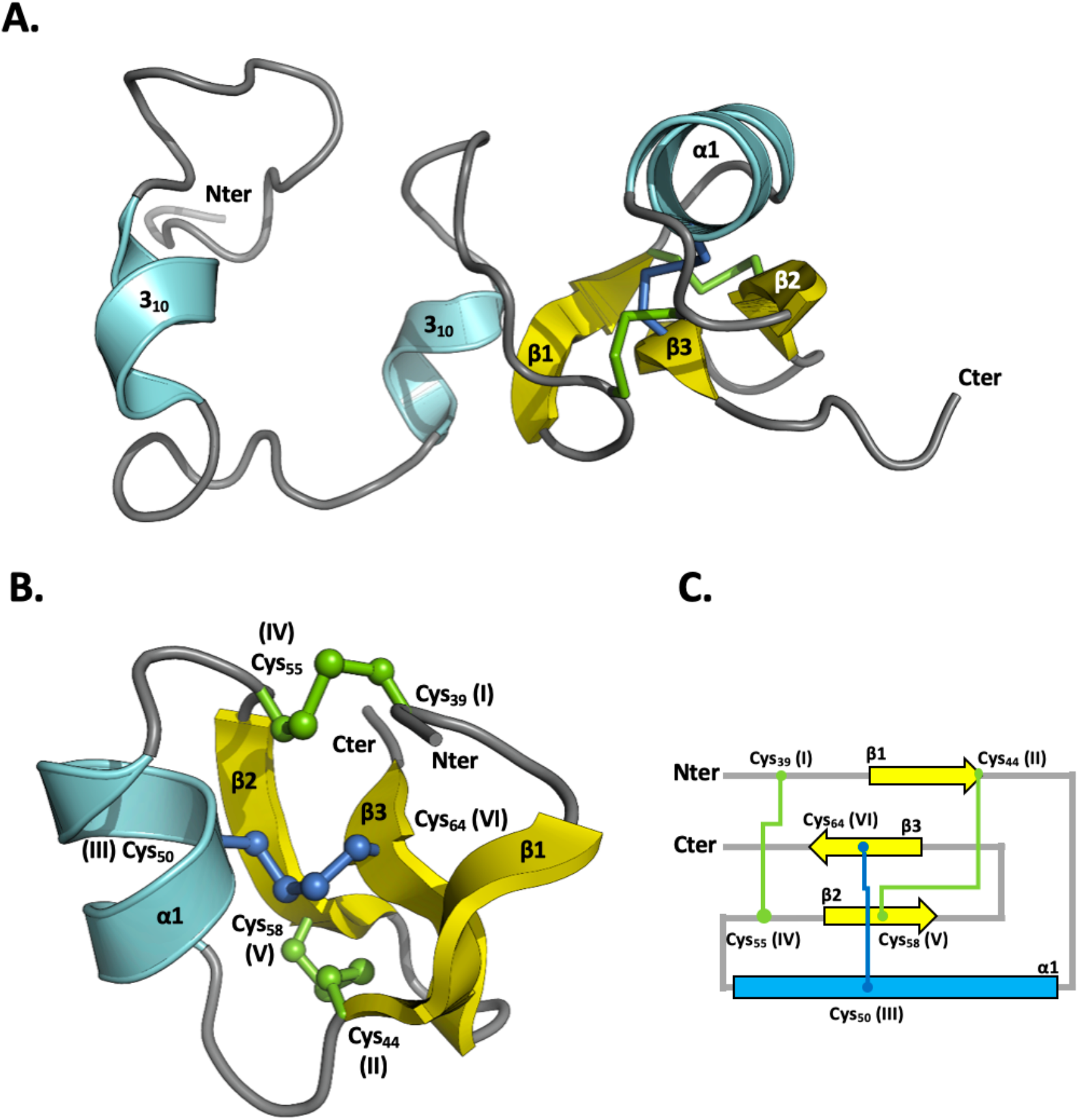
NMR spectroscopy solution structure of MLP124266. **A.** General structure of MLP124266 represented as cartoon. **B.** The C-terminal small disulfide-rich domain is related to the structural family of knottins, which contain at least 3 disulfide bridges and 6 cysteines and implies a disulfide between cysteines III and VI going through disulfides I-IV and II-V. In MLP124266, a disulfide bridge between Cys50 (III) and Cys64 (VI), in blue, goes through disulfide bridges between Cys39 (I) and Cys55 (IV) as well as Cys44 (II) and Cys58 (V), in green. **C.** Schematic representation of disulfide bridge connectivity. Disulfide bridge Cys50 (III)-Cys64 (VI) is represented in blue and disulfide bridges between Cys39 (I)-Cys55 (IV) and Cys44 (II)-Cys58 (V) are represented in green. Helix α (residues 47 to 53) is coloured in cyan.

### MLP124017 is part of the Nuclear Transport Factor 2-like protein superfamily

MLP124017 is a small-secreted protein (167 amino acids with its signal peptide; 150 amino acids in its mature form, with a molecular mass of 18 kDa) of unknown function, highly expressed during infection of poplar leaves by *M. larici-populina* (Hacquard *et al.*, 2012). MLP124017 shares sequence similarity with neither other *M. larici-populina* nor other rust fungal proteins. In a previous study, we demonstrated the nucleocytoplasmic localization of MLP124107 in *N. benthamiana* and its interaction with poplar TOPLESS-related 4 protein (Petre *et al.*, 2015). To further investigate MLP124017 structure and to get insights about its function, we first attempted to solve its 3D structure by crystallization coupled to X-ray diffraction. We were unable to obtain exploitable diffracting crystals despite the use of different versions (untagged or N- or C-terminal His-tagged) of MLP124017 protein and therefore switched to NMR. The recombinant ^15^N and ^13^C-labelled MLP124017 protein was used for structure determination by two- and three-dimensional NMR experiments (Table S3). The assigned ^1^H, ^15^N-HSQC spectra were well dispersed but the peaks for residues from the N-terminal 1-14 and 86-95 segments were missing (Figure S5). From preliminary structures, the production of a truncated recombinant protein for the first eight N-terminal residues that could mask residues 86-95 did not improve the data. The solution structure of MLP124017 was determined based on 1727 NOE-derived distance restraints, 214 dihedral angle restraints and 102 hydrogen bond restraints. All proline residues have been determined to be in a trans-conformation according to the ^13^Cβ chemical shift at 32.21; 32.46; 31.02 and 32.40 ppm for Pro_36_; Pro_51_; Pro_54_ and Pro_146_ respectively. The best conformers with the lowest energies, which exhibited no obvious NOE violations and no dihedral violations > 5° were selected for final analysis. The Ramachandran plot produced shows that 99.6% of the residues are in favoured regions (Table S4). MLP124017 structure is composed of a α+β barrel with seven β-strands forming one mixed β-sheet, four β-hairpins, four β-bulges, and four α-helices (Figure 5A). Residues 1-14 and 150-151 having missing assignments are not defined in the final models. This arrangement of secondary structure produces a cone-shaped fold for the protein, which generates a distinctive hydrophobic cavity (Figure 5B). To identify potential structural homologs of MLP124017, we performed structural similarity searches using the Dali server (Holm and Rosenström, 2010). Queries identified SBAL_0622 (PDB code 3BLZ) and SPO1084 (PDB code 3FKA) as the closest structural homologs with the highest Z-score of 6.0 and 6.3, respectively, and a RMSD of 4.1 and 3.5Å, respectively (Figure S6). These two proteins, which are from the bacteria *Shewanella baltica* and *Ruegeria pomeroyi*, have no known function, but share a common Nuclear Transport Factor 2-like (NTF2) fold. The NTF2 superfamily comprises a large group of proteins that share a common fold and that are widespread in both prokaryotes and eukaryotes (Eberhardt *et al.*, 2013). Taken together these results show that although MLP124017 do not share sequence similarities or domain with other proteins in sequence databases, its structure is similar to proteins of the NTF2 folding superfamily.

**Figure 5:**
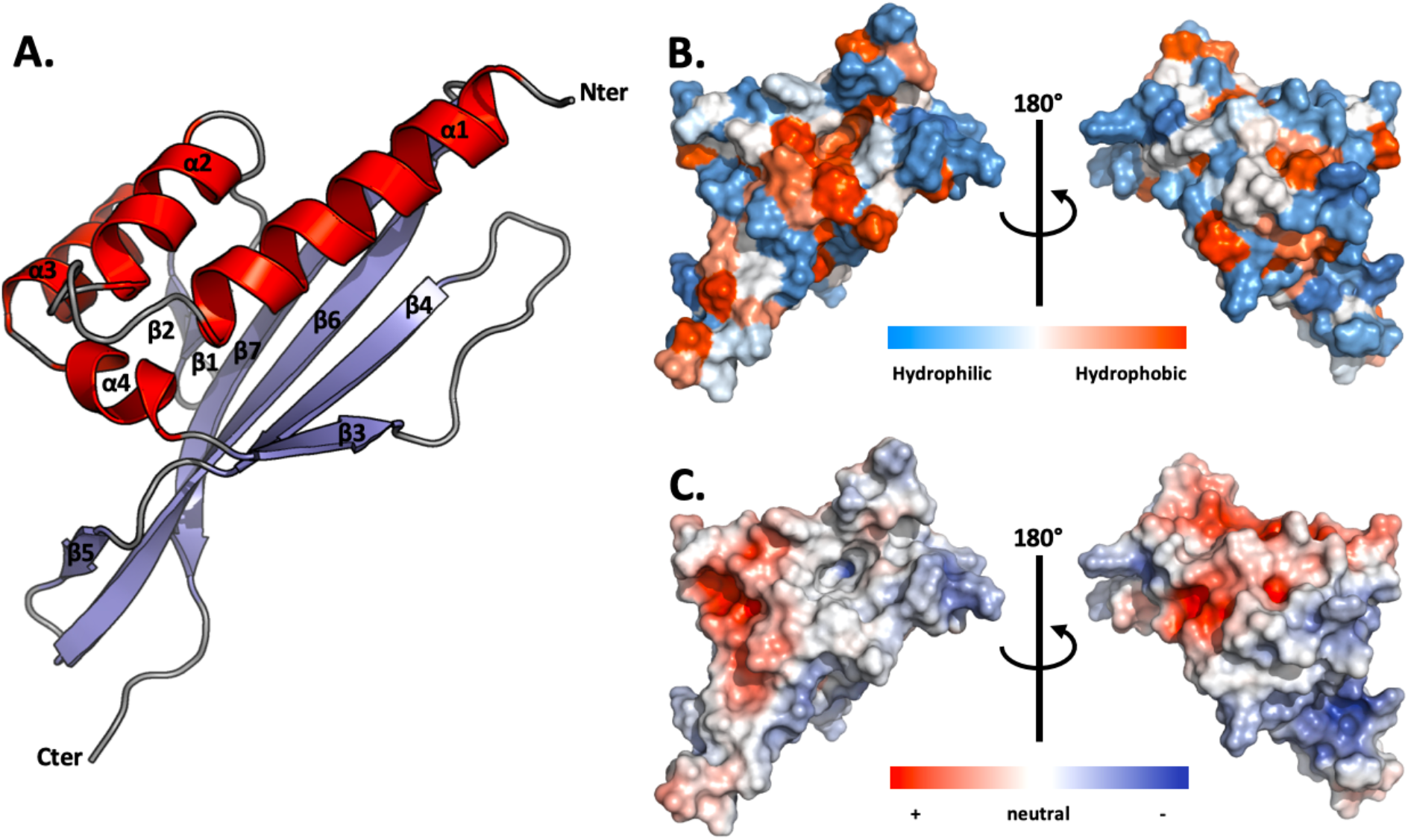
NMR spectroscopy solution structure of MLP124017. **A.** The structure of MLP124017 is represented as cartoon and consists of four α-helices in red (α1: residues 16-32; α2: 41-48; α3: 65-73; α4: 80-82) and a mixed β-sheet composed by seven β-strands coloured in blue (β1: residues 55-58; β2: 61-63; β3: 84-86; β4: 98-104; β5: 108-109; β6: 115-129; β7: 132-144). **B.** Front and rear views on the surface of MLP124017 illustrating the surface hydrophobic potential. The hydrophobic and hydrophilic patches are shown in orange and in blue respectively. **C.** Front and rear views on the surface of MLP124017 illustrating the surface electrostatic Coulomb potential at pH 7.0 using APBS plugin from Pymol 2.3 software with a contour of −10 kT/e to 10 kT/e. The positive-charge and negative-charge densities are coloured in blue and red respectively.

## DISCUSSION

In this study, we have set up a small-throughput effectoromics pipeline based on recombinant protein production and structural characterization to get insights on eleven candidate effectors of the poplar leaf rust fungus *M. larici-populina*. Out of the eleven selected effectors, three were successfully purified as recombinant proteins, and two were structurally characterized by NMR spectroscopy. MLP124266 is a homolog of the *M. lini* AvrP4 effector protein (Catanzariti *et al.*, 2006), and we showed that it exhibits a cystine-knot (or knottin) structural motif commonly encountered in small disulfide-rich proteins. MLP124017 is an orphan protein in *M. larici-populina* with no known ortholog in Pucciniales. MLP124017 physically associates with the poplar TOPLESS-related protein 4 (TRP4) (Petre *et al.*, 2015), and we showed that it exhibits a fold similar to two bacterial proteins that belong to the Nuclear-Transport factor 2-like protein superfamily.

Recombinant protein production in *E. coli* is a valuable approach to perform biophysical and biochemical analyses of candidate effectors (Franceschetti *et al.*, 2017; Zhang *et al.*, 2017). However, we have faced issues for the production of soluble small-cysteine rich proteins in this system. Indeed, among the eleven candidate effectors screened for expression only three were found in the soluble protein fraction and stable enough along the purification procedure. Although we tested different *E. coli* strains and protein expression induction protocols, the eight other candidate effectors were either not expressed at all or expressed as inclusion bodies. It is possible to purify recombinant proteins from inclusion bodies by using denaturing extraction conditions and further refolding proteins (Palmer and Wingfield, 2012). However, this approach is not recommended for structural analysis as the refolding of the proteins *in vitro* may alter folding. Another limit of prokaryote systems to produce eukaryote proteins is the lack of post-translational modifications such as methylations. An alternative to this is to use the yeast *Pichia pastoris*, which has proven useful for several fungal effectors such as *Leptospheria maculans* AvrLm4-7 or *Cladosporium fulvum* Avr2 and Avr4 (Blondeau *et al.*, 2015; Rooney *et al.*, 2005). We have tried this system to produce the candidate effector (MLP107772), but without success. Nevertheless, this system may be useful and deserves to be considered as an alternative to assay other rust effectors for which we were not able to obtain production in *E. coli*.

We showed that MLP124266 possesses two distinct regions with contrasted structural properties. The C-terminal part is rigidified by a cystine-knot motif, whereas the N-terminal part is globally flexible. The knottin folded proteins display a variety of functions such as venoms and spider toxins (Garcia, 2004; Lee and MacKinnon, 2004) but also antimicrobial properties such as the cyclotides (Tam *et al.*, 1999). Some are also found to interact with receptor like protease inhibitors found in plants, insects and plant parasites (Kim *et al.*, 2009). The three disulfide bridges within the C-terminal part of MLP124266 confer its rigidity, as confirmed by our NMR dynamic study, and probably contribute to the high protein stability (Daly *et al.*, 2011). MLP124266 presents a β-sheet structure typical of knottins, but interestingly it also has an additional helix between β2 and β3 strands. To our knowledge, the presence of such a helix in knottins has been reported in cyclotides only, and more precisely in bracelet cyclotides containing six or seven residues in the loop between Cys(III) and Cys(IV) (Chen *et al*, 2005). This loop often contains an alanine, which favours the formation of the helix as well as a highly conserved glycine allowing its connection to the cystine knot (Rosengren *et al.*, 2003). Interestingly, the loop in MLP124266 has such residues, *i.e.* Ala^52^ and Gly^54^, but consists of four residues only. In *Viola odorata* cycloviolacin O2 (cO2), the additional helix is located in a hydrophobic loop that interacts with the membrane-mimicking micelles (Wang *et al.*, 2009). Therefore, it might help disrupting membranes and thus contribute to the cytotoxicity activity of cO2 and play a role in plant defence. In MLP124266, the helical turn is not particularly hydrophobic (Figure S4B) and may not have these properties. To our knowledge, MLP124266 is the first fungal protein to present a knottin-like structure (Postic *et al.*, 2018). It would be interesting to collect structural data from other potential fungal knottins to find out whether the additional helix is always present and to clarify its role.

Several properties of MLP124266 N-terminal region also deserve to be pointed out. This region, approximately extending up to residue 35 and thus representing half of the primary sequence, globally presents a higher flexibility, as demonstrated by the NMR dynamic results. Nevertheless, it is not totally unstructured, as the heteronuclear NOE values remain positive and as a few residues exhibit gamma turn propensity. This results in the presence of short helical turns, especially in the segment 8-17 and to a lower extent for residues 28-30. Although these secondary structures are not systematically found in all NMR models and their limits are very variable, several secondary structure prediction softwares predict a helix between residues 11 and 18. Moreover, the proportion of conserved amino acids is strikingly high in this N-terminal region, much higher than in the folded C-terminal part (Figure S4). Altogether, these results suggest that the relatively flexible N-terminal region have some propensity to form helical structures, which may support a biochemical role that remains to be elucidated. Interestingly, other effectors possess a predicted disordered N-terminal region (Boutemy *et al.*, 2011). For instance, *M. lini* effectors AvrL567 and AvrM have predicted disordered N-terminal regions that are susceptible to protease degradation (Catanzariti *et al.*, 2006; Wang *et al.*, 2007). Flexible folds are known to be adaptable linkers that favour the ability to bind partners (Chouard, 2011). As the N-terminus of many cytoplasmic effectors is anticipated to mediate protein entry into host cells (Petre and Kamoun, 2014), it is tempting to speculate that this flexible part may bind a target important for cell entry. MLP124266 belongs to a rust multigene family reported in the Melampsoraceae for which population data has been collected. The survey of synonymous and non-synonymous nucleotide substitutions in isolates of *M. lini*, *M. larici-populina* and in several *Melampsora* spp. identified positions under positive selection in between strictly conserved cysteines in the C-terminus region of the proteins (Barrett *et al.*, 2009; Hacquard *et al.*, 2012; Persoons *et al.*, 2014; Van der Merwe *et al.*, 2009). In the flax rust fungus *M. lini*, the homolog of MLP124266 is an avirulence protein that evolved under the pressure of the plant host immune system. The resolution of the MLP124266 (AvrP4 homolog) structure suggests that its compact and structured C-terminal region may be more prone to detection by plant immune receptors.

We solved the structure of MLP124017 by NMR spectroscopy. MLP124017 showed a fold similar to members of the NTF2-like superfamily. The NTF2-like superfamily is a group of protein domains sharing a common fold, but showing no sequence similarity. MLP124017 is structurally similar to two bacterial proteins, despite the lack of sequence similarity. The structures of these two bacterial proteins consist of a β-sheet surrounding a binding pocket and α-helices acting as a lid (Marcos *et al.*, 2017). The NTF2 family regroups catalytic and non-catalytic proteins that contain cone-like structured proteins with a cavity that often act as a molecular container involved in a wide range of cellular functions (Eberhardt *et al.*, 2013). Interestingly, the cone-shaped structure of MLP124017 is widespread across both prokaryotes and eukaryotes. The first proteins of the NTF2 family were reported to play a role in the transport of molecules from the cytoplasm to the nucleus. Arabidopsis NTF2 protein is required to import nuclear proteins via the recognition of a nuclear localization signal (NLS). This protein also plays a role in the nuclear import of the small-GTPase Ran-GDP that is a central protein in various signal transduction pathways (e.g. mitotic spindle formation, nuclear envelope assembly, or responses to biotic stresses; (Avis and Clarke, 1996; Carazo-Salas *et al.*, 2001; Hetzer *et al.*, 2000; Zhang and Clarke, 2000). In bacteria, some NTF2-like proteins play a role in bacterial conjugation as part of the type IV secretion system (Goessweiner-Mohr *et al.*, 2013), whereas non-catalytic NTF2-like domains act as immunity proteins (Zhang *et al.*, 2012). In fungi, *Saccharomyces cerevisiae* NTF2 mutant is defective for nuclear import (Corbett and Silver, 1996). Although NTF2 folded proteins are widespread across kingdoms, very few is known about their role. A recent study presented that the silencing of NTF2 in wheat decreased the resistance against avirulent isolates of the wheat stripe rust fungus *P. striiformis* f. sp. *Tritici* (Zhang *et al.*, 2018). Since MLP124017 has been shown to interact with TOPLESS and TOPLESS-related proteins (Petre *et al.*, 2015), it is tempting to speculate that the cavity formed by the β-sheet could be involved in the association with these plant partners.

Although MLP124266 and MLP124017 show no primary sequence similarity to known proteins, they adopt a three dimensional fold similar to knottins or NTF2 family members. Thus, knowing the structure of both candidate effectors allowed us to classify them as members of large superfamilies of proteins. The concept of structural families whose members share no, or very limited, primary sequence homology emerges in effector biology (Białas *et al.*, 2018). This concept promises to revolution the way we predict and categorize effector proteins (de Guillen *et al.*, 2015). For instance, members of the MAX effector family share a common β-sandwich fold, but show no primary sequence similarity (de Guillen *et al.*, 2015). Similarly, members of the WY superfamily of RXLR effectors in oomycetes share a common three-to four-helix bundle (Boutemy *et al.*, 2011). Such features are now used to search and categorize fungal and oomycete effector proteins into structural superfamilies (de Guillen *et al.*, 2015; Win *et al.*, 2012). To our knowledge, MLP124266 and MLP124017 are the first effector proteins to adopt knottins and NTF2 folds. Whether other effector proteins adopt similar folds remains to be identified to determine if knottins and NTF2 folds define structural superfamilies in fungi.

To conclude, in this study we screened eleven rust candidate effectors using biochemical and structural approaches, and managed to solve the structure for two of them. This demonstrates the usefulness of such approaches for systematically screening candidate effectors and gaining information about their putative biological roles. Although it is too early to speculate about the function of MLP124266 and MLP124017, these structures form a basis for future comparative structural studies and bring complementary information regarding fungal knottins.

## EXPERIMENTAL PROCEDURES

### Sequence analyses and names

Alignment and phylogenetic analyses were performed on the phylogeny website (www.phylogeny.fr) with default parameters. Alignments were corrected and edited manually, and phylogenetic trees were generated with FigTree v1.4.3 (http://tree.bio.ed.ac.uk/software/figtree/). Physical and chemical parameters of proteins were estimated using protparam tool (http://web.expasy.org/protparam/). Common names and JGI ID of genes described in this study are as follow: *Mlp124266*, *Mlp124499*, *Mlp124111*, *Mlp124478*, *Mlp124530*, *Mlp124561*, *Mlp37347*, *Mlp109567*, *Mlp124017*, *Mlp107772*,and *Mlp124202*. The mapping of the family-wide conservation pattern of amino acids onto MLP124266 structure was performed with Consurf (http://consurf.tau.ac.il/2016/).

### Cloning of selected effector encoding sequences

Open reading frames coding for the mature forms (i.e. devoid of the sequence encoding N-terminal secretion peptide) of MLP124266 and MLP124499 of *M. larici-populina* isolate 98AG31 were ordered as synthetic genes cloned in pBSK(+) vectors (Genecust). Coding sequence of the mature forms of the nine other candidate effectors (*Mlp124111*, *Mlp124478*, *Mlp124530*, *Mlp124561*, *Mlp37347*, *Mlp109567*, *Mlp124017*, *Mlp107772*, and *Mlp124202*) were amplified by polymerase chain reaction (PCR) using cDNAs from leaves of the poplar hybrid Beaupré infected by *M. larici-populina* (isolate 98AG31) and further cloned into pICSL01005 vector as described previously (Petre *et al.*, 2015). The sequences encoding the mature form of each effector were subsequently cloned by PCR in either pET26b or pET28a vector between *Nde*I and *Not*I or *Nco*I and *Not*I restriction sites, respectively, using primers shown in Table S1.

### Expression and purification of recombinant proteins in *Escherichia coli*

Expression of recombinants proteins was performed at 37°C using the *E. coli* SoluBL21 (DE3) pRARE2 (Amsbio Abington, UK), Rosetta2 (DE3) pLysS, Origami2 (DE3) pLysS or RosettaGami2 (DE3) pLysS strains (Novagen) containing the adequate pET expression vector coding for the selected candidate effector (Table S1) in LB medium supplemented with kanamycin (50 μg/ml) and chloramphenicol (34 μg/ml). When the cell culture reached an OD_600_nm of 0.7, protein expression was induced with 0.1 mM isopropyl-β-D-1-thiogalactopyranoside (IPTG) and cells were grown for a further 4 h. To improve the solubility of some recombinant candidate effectors, other protocols were used as follows. First, we added 0.5% (v/v) of ethanol in the medium when culture reached an OD_600_nm of 0.7. The cells were cooled to 4°C for 3 h, recombinant protein expression induced with 0.1 mM IPTG and cells further grown for 18 h at 20°C. We also tested a combination of an osmotic and a thermal shock (Oganesyan *et al.*, 2007). When the culture reached an OD_600_nm of 0.5-0.6, 500 mM NaCl and 2 mM of betaine were added to the culture medium and the culture incubated at 47°C for 1 hour under stirring. Cells were then cooled to 20°C and the expression of recombinant proteins induced with 0.1 mM IPTG. After induction, cells were harvested by centrifugation, suspended in a 30 mM Tris-HCl pH 8.0 and 200 mM NaCl lysis buffer, and stored at −20°C. Cell lysis was completed by sonication (three times for 1 min with intervals of 1 min). The cell extract was then centrifuged at 35 000 g for 25 min. at 4 °C to remove cellular debris and aggregated proteins. C-terminal His-tagged recombinant proteins were then purified by gravity-flow chromatography on a nickel nitrilotriacetate (Ni-NTA) agarose resin (Qiagen) according to the manufacturer’s recommendations followed by an exclusion chromatography on a Superdex75 column connected to an ÄKTA Purifier^TM^ (GE Healthcare). The fractions containing recombinant MLP124017 were pooled, concentrated, and stored at −20°C as such, whereas for MLP124266 and MLP124499 fractions were pooled, dialyzed against 30 mM Tris-HCl, 1 mM EDTA (TE) pH 8.0 buffer, and stored at 4°C until further use.

For the NMR spectroscopy analyses, MLP124266-(His6) and MLP124499-(His6) recombinant proteins were ^15^N-labeled in M9 minimal synthetic medium containing ^15^NH_4_Cl (1 g/L). MLP124017 was single ^15^N or double ^15^N and ^13^C labelled in M9 minimal medium containing 1 g/l NH_4_Cl (^15^N) and 2 g/l glucose (^13^C) supplemented with 2,5% thiamine (m/v), 1mg/ml biotin, 50 mM FeCl_3_, 10 mM MnCl_2_, 10 mM ZnSO_4_, 2 mM CoCl_2_, 2 mM NiCl_2_, 2 mM NaSeO_3_, and 2 mM H_3_BO_3_. After purification as described above, labelled MLP124266, MLP124499, and MLP124017 were dialyzed against a 50 mM phosphate pH 6.0 or a 20 mM phosphate pH 6.8 buffer supplemented with 200 mM NaCl. The homogeneity of purified proteins was checked by SDS-PAGE and protein concentration determined by measuring the absorbance at 280 nm and using theoretical molar absorption coefficients of 500 M^−1^.cm^−1^, 3 105 M^−1^.cm^−1^, 29 450 M^−1^.cm^−1^ deduced from the primary sequences of mature MLP124266, MLP124499, and MLP124017 proteins respectively. For MLP124266, protein concentration was also verified using a colorimetric assay (BC assay, Interchim).

### Protein sample preparation for NMR spectroscopy

Uniformly labelled ^15^N MLP124017 (1 mM in 20 mM phosphate pH 6.8, 200 mM NaCl) was supplemented with 0.02% sodium azide and 10% of 5 mM 4,4-dimethyl-4-silapentane-1-sulfonic acid (DSS) in D_2_O as a lock/reference. For the D_2_O experiments, the sample was lyophilized and dissolved in 200 µL D_2_O. For the 3D heteronuclear experiments, a ^13^C/^15^N labelled sample at 0.6 mM was prepared with 200 µL in the same phosphate buffer previously used supplemented with 20 µL of D_2_O at 5 mM DSS as a reference. MLP124266 and MLP124499 samples were dissolved in 50 mM phosphate pH 6.0 buffer with 10% D_2_O and 0.02% sodium azide. The concentration of unlabelled MLP124266 and MLP124499 was 0.85 and 0.65 mM, and for uniformly ^15^N labelled MLP124266 and MLP124499 was 1.8 and 0.135 mM, respectively.

### Nuclear magnetic resonance spectroscopy

For MLP124017, spectra were acquired on 800 and 700 MHz Avance Bruker spectrometers equipped with triple-resonance (^1^H, ^15^N, ^13^C) z-gradient cryo-probe at 298 K. Experiments were recorded using the TOPSPIN pulse sequence library (v. 2.1) (Table S2). All spectra are referenced to the internal reference DSS (4,4-dimethyl-4-silapentane-1-sulfonic acid) for the ^1^H dimension and indirectly referenced for the ^15^N and ^13^C dimensions (Wishart *et al.*, 1995). Sequential assignment was performed using 3D ^15^N-NOESY-HSQC, ^15^N-TOCSY-HSQC, HNCO, HNCACO, HNCA, HNCOCACB, and HNCACB. Side chain ^1^H assignments were carried out using combined analysis with 3D ^15^N-NOESY-HSQC, ^15^N-TOCSY-HSQC, and 2D NOESY and TOCSY with D_2_O samples. A series of three HSQC spectra were performed after lyophilisation and dilution of the first sample in D_2_O to determine amide protons in slow exchange (Table S2).

For MLP124266 and MLP1124499, NMR spectra were acquired on a Bruker DRX 600 MHz spectrometer equipped with a TCI cryoprobe. For MLP124266 and MLP124499, COSY, TOCSY (mixing time of 60 ms) and NOESY (mixing time of 150 ms) experiments were run at 298 K, respectively. For MLP124266, HNHA, HNHB, R1 and R2 ^15^N relaxation rates, ^1^H-^15^N heteronuclear NOE, HNCA (with 24 (^15^N) × 28 (^13^C) complex points and 192 transients per increment) standard experiments were recorded. Spectra were processed using Topspin® 3.0 software (Bruker) and analysed with NmrViewJ (Johnson and Blevins, 1994), CcpNmr (Vranken *et al.*, 2005) and ARIA2 (Rieping *et al.*, 2007).

### Structure calculation

NOE peaks identified in 3D ^15^N-NOESY-HSQC and 2D NOESY experiments were automatically assigned during structure calculations performed by the program CYANA 2.1 (Güntert, 2004). The ^15^N, H_N_, ^13^C’, ^13^Cα, Hα, and ^13^Cβ chemical shifts were converted into φ/Ψ dihedral angle constraints using TALOS+ (v. 1.2) (Shen *et al.*, 2009). Hydrogen bonds constraints were determined according to ^1^H/^2^H exchange experiments of backbone amide protons (H_N_). Each hydrogen bond was forced using following constraints: 1.8–2.0 Å for H_N_,O distance and 2.7–3.0 Å for N_H_,O distance. Final structure calculations were performed with CYANA (v. 2.1) using all distance and angle restraints (Table S3). 600 structures were calculated with CYANA 2.1, of which the 20 conformers with the lowest target function were refined by CNS (v. 1.2) using the refinement in water of RECOORD (Nederveen *et al.*, 2005) and validated using PROCHECK (Laskowski *et al.*, 1993).

For MLP124266 structure calculation, 783 NOE peaks identified in 3D ^15^N-NOESY-HSQC or 2D NOESY spectra, 75 φ/Ψ dihedral angles generated by DANGLE (Cheung et al, 2010) and 10 hydrogen bonds were input as unambiguous restraints in ARIA2. Covalent disulfide bonds between Cys39-Cys55, Cys44-Cys58 and Cys50-Cys64 were also introduced. Among the 400 structures generated by ARIA2, 20 models of lowest energy were refined in water (Table 2).

NMR assignment and structure coordinates have been deposited in the Biological Magnetic Resonance Data Bank (BMRB code 34298 and 34423) and in the RCSB Protein Data Bank (PDB code 6H0I and 6SGO), respectively.

## ACKNOWLEDGMENTS

The UMR1136 is supported by a grant overseen by the French National Research Agency (ANR) as part of the “Investissements d’Avenir” program (ANR-11-LABX-0002-01, Lab of Excellence ARBRE). This work was supported by the French Infrastructure for Integrated Structural Biology (ANR-10-INBS-0005).

## DATRA AVAILABILITY STATEMENT

The data that support the findings of this study are openly available in the Biological Magnetic Resonance Data Bank (http://www.bmrb.wisc.edu/) and in the RCSB Protein Data Bank (http://www.rcsb.org/).

## AUTHOR’S CONTRIBUTION

AH, NR and SD contributed to the design and implementation of the research. BP, CL, KG, NS, PT performed the experiments. KG, PB and PT performed structural data analyses. All authors participated to the analysis of the results and to the writing of the manuscript.

---

**Table S1:**
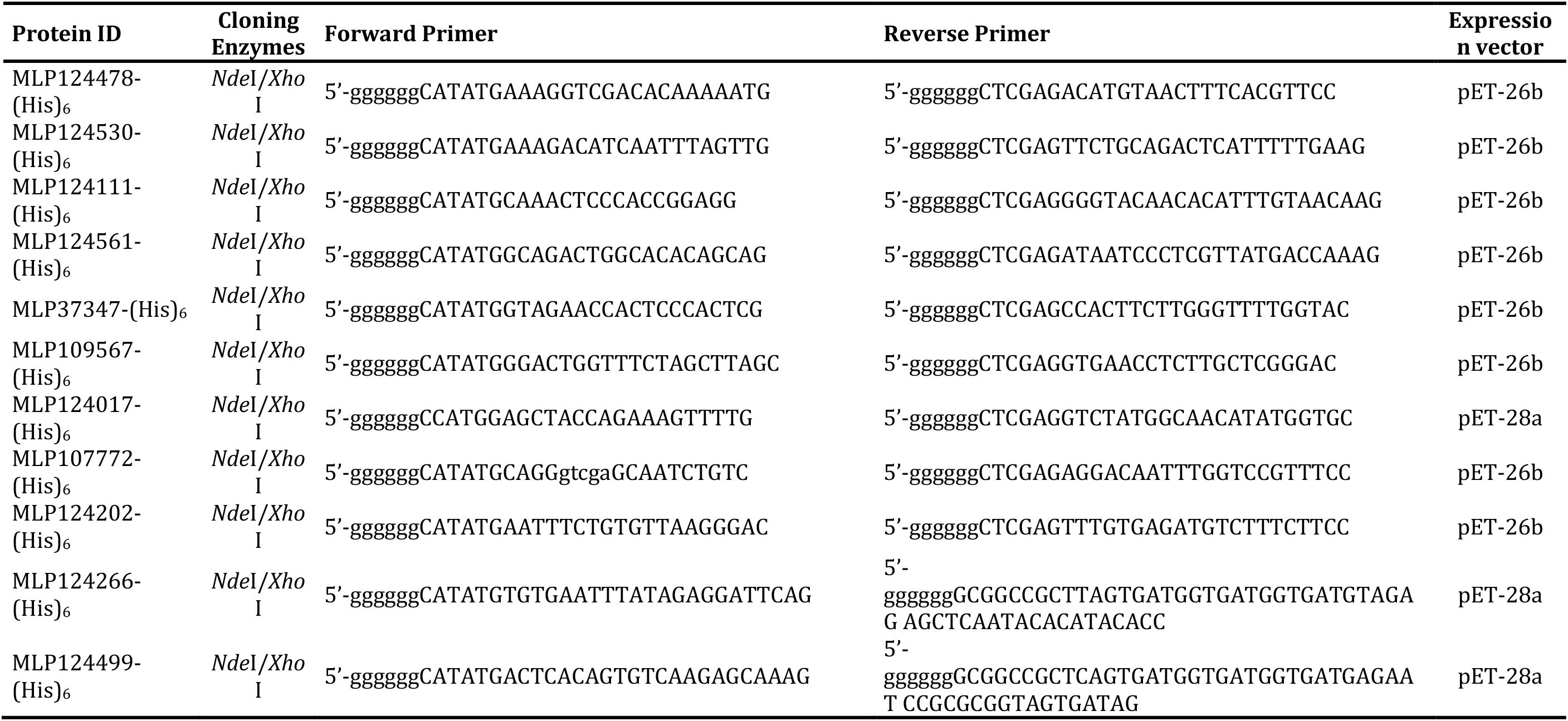
Primers used for PCR cloning in pET expression vectors

**Table S2:** Statistics for 20 NMR structures of MLP124266

|  | MLP124266 |
| --- | --- |
| NOE restraints | 653 |
| Short range ($\vert i-j\vert \leq 1$) | 510 |
| Medium range ($1 < \vert i-j\vert < 5$) | 102 |
| Long range ($\vert i-j\vert \geq 5$) | 41 |
| H-bond restraints | 7 |
| Dihedral restraints <sup>(a)</sup> | 75 |
| Number of NOE violations |  |
| > 0.0 Å | 82 |
| > 0.3 Å | 45 |
| > 0.5 Å | 18 |
| Dihedral violations |  |
| > 0° | 63 |
| > 5° | 23 |
| Ramachandran plot statistics |  |
| most favourable regions (%) | 81.9 |
| allowed regions (%) | 15.3 |
| disallowed regions (%) | 2.8 |
| RMSD (Å) <sup>(b)</sup> |  |
| Backbone | 1.44 ± 0.52 |
| Heavy atoms | 2.08 ± 0.53 |

<sup>(a)</sup> Dihedral restraints were generated using DANGLE.
<sup>(b)</sup> RMSD were calculated over residues 39 to 69.

**Table S3:** NMR experiments acquired for structure calculations and chemical shift assignments for MLP124017

| Experiments | nuclei | Size |  |  | Sweep width (ppm) |  |  | Mix(ms) | NS | D1(s) | B<sub>0</sub>(MHz) |
| --- | --- | --- | --- | --- | --- | --- | --- | --- | --- | --- | --- |
|  |  | F3 | F2 | F1 | F3 | F2 | F1 |  |  |  |  |
| <sup>15</sup>N-HSQC | <sup>1</sup>H, <sup>15</sup>N | 1800 | 180 | - | 13.94 | 36 | - | - | 4 | 1.0 | 800 |
| HNCO(*) | <sup>1</sup>H,<sup>15</sup>N,<sup>13</sup>C | 1366 | 60 | 100 | 13.94 | 36 | 15 | - | 8 | 0.2 | 800 |
| HNCA(*) | <sup>1</sup>H,<sup>15</sup>N,<sup>13</sup>C | 1362 | 50 | 60 | 13.94 | 36 | 30 | - | 16 | 0.2 | 800 |
| HNCOCACB(*) | <sup>1</sup>H,<sup>15</sup>N,<sup>13</sup>C | 1366 | 60 | 100 | 13.94 | 36 | 70 | - | 32 | 0.2 | 800 |
| HNCACO(**) | <sup>1</sup>H,<sup>15</sup>N,<sup>13</sup>C | 1366 | 40 | 100 | 13.94 | 36 | 15 | - | 16 | 1.0 | 800 |
| HNCACB(*) | <sup>1</sup>H,<sup>15</sup>N,<sup>13</sup>C | 1362 | 50 | 60 | 13.94 | 36 | 60 | - | 32 | 0.2 | 800 |
| <sup>15</sup>N-NOESY-HSQC | <sup>1</sup>H,<sup>15</sup>N, <sup>1</sup>H | 1800 | 64 | 360 | 13.94 | 25 | 13.94 | 150 | 8 | 1 | 800 |
| <sup>15</sup>N-TOCSY-HSQC | <sup>1</sup>H,<sup>15</sup>N, <sup>1</sup>H | 1800 | 64 | 360 | 13.94 | 25 | 13.94 | 54 | 8 | 1 | 800 |
| NOESY (D<sub>2</sub>O) | <sup>1</sup>H, <sup>1</sup>H | 2048 | 512 | - | 12.03 | 12.03 | - | 150 | 128 | 1 | 700 |
| TOCSY (D<sub>2</sub>O) | <sup>1</sup>H, <sup>1</sup>H | 2048 | 512 | - | 12.03 | 12.03 | - | 57.6 | 128 | 1 | 700 |
| <sup>15</sup>N-HSQC (***) | <sup>1</sup>H, <sup>15</sup>N | 1112 | 256 | - | 15.95 | 36 | - | - | 2 | 0.2 | 700 |

Experiments were recorded using the TOPSPIN Library (v. 2.1) at 298 K.
(\*) BEST pulse sequences
(\*\*) Watergate version
(\*\*\*) SOFAST version (from Institut de Biologie Structurale, Grenoble,
http://www.ibs.fr/research/scientific-output/software/pulse-sequence-tools/article/ibs-pulse-sequence-tools-for-bruker-spectrometers)

**Table S4:** Statistics for 20 NMR structures of MLP124017

|  | MLP124017 |
| --- | --- |
| NOE restraints | 1727 |
| Short range ($\vert i-j\vert \leq 1$) | 999 |
| Medium range ($1 < \vert i-j\vert < 5$) | 345 |
| Long range ($\vert i-j\vert \geq 5$) | 383 |
| H-bond restraints | 102 |
| Dihedral restraints <sup>(a)</sup> | 214 |
| Number of NOE violations |  |
| $> 0.0 \text{ \AA}$ | 186.35 |
| $> 0.3 \text{ \AA}$ | 0.05 |
| $> 0.5 \text{ \AA}$ | 0 |
| Dihedral violations |  |
| $> 0^\circ$ | 22.25 |
| $> 5^\circ$ | 0 |
| Ramachandran plot statistics |  |
| Most favourable regions (%) | 89.8 |
| Additionally allowed regions (%) | 9.8 |
| Generously allowed regions (%) | 0.4 |
| Disallowed regions (%) | 0.1 |
| RMSD ($\text{\AA}$) <sup>(b)</sup> | |
| Backbone | $1.24 \pm 0.39$ |
| Heavy atoms | $1.81 \pm 0.40$ |

<sup>(a)</sup> Residues in regular secondary structures were derived from the chemical shifts using TALOS+ software.
<sup>(b)</sup> Main chain atoms (N, Cα , C) over the residues 20-81 and 100-141.

**Figure S1:**
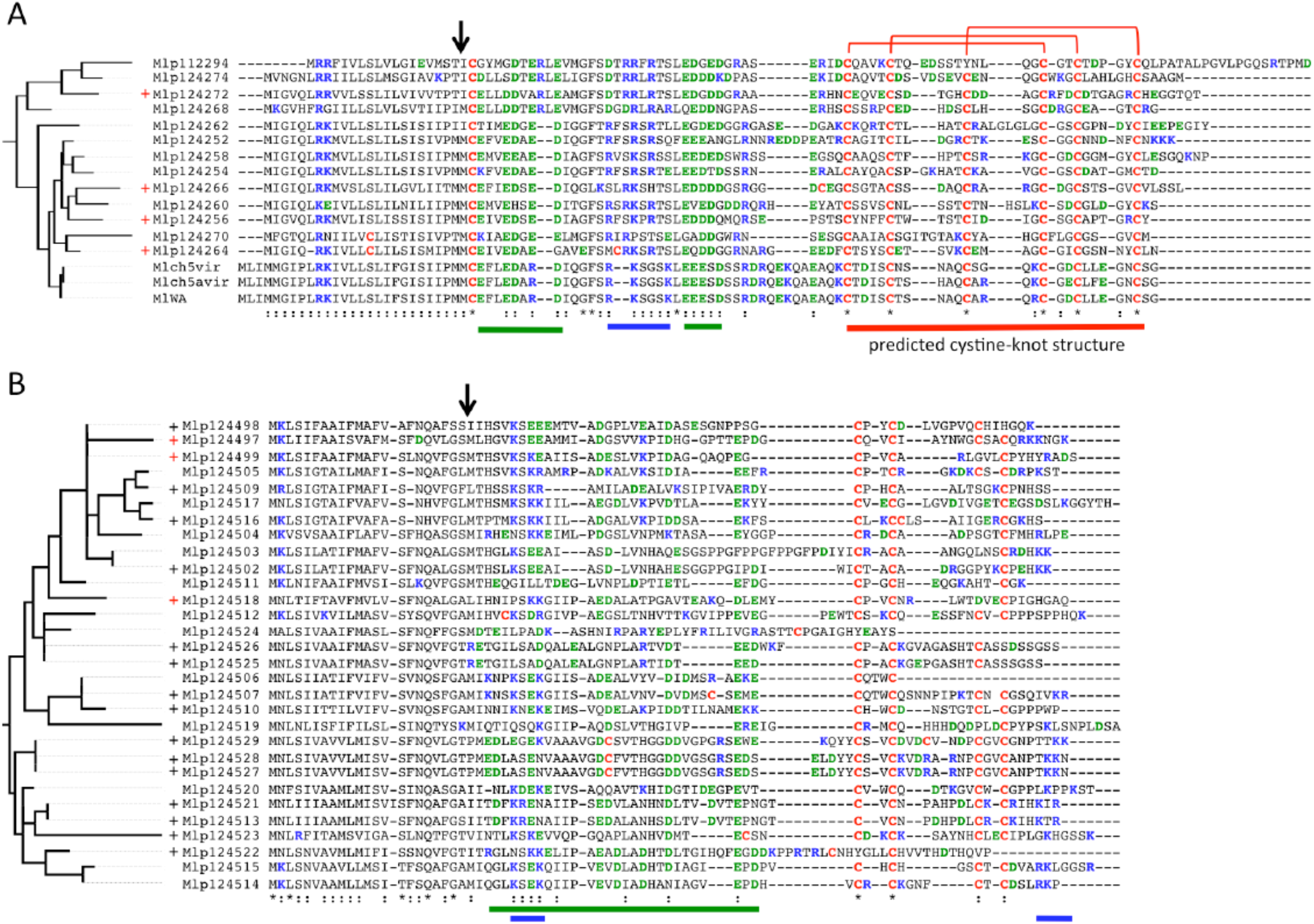
Sequence characteristics of proteins from MLP124266 and MLP124499 families. Phylogenetic trees were built from corresponding aligned protein sequences from MLP124266 (**A**) and MLP124499 (**B**) families. Branch length is proportional to phylogenetic distance. Black ‘+’ indicates that expression of the gene has been detected *in planta* (Duplessis *et al.*, 2011; Joly *et al.*, 2010), red ‘+’ indicates that the RT-qPCR expression profile of the gene has been established (Hacquard *et al.*, 2012). The black arrows indicate the predicted cleavage site of the signal peptide. Cysteine residues are in red, basic residues (lysine/K and arginine/R) are in blue, acid residues (aspartic acid/D and glutamic acid/E) are in green. Blue and green lines show basic and acid stretches of amino acids and the red line delimited the predicted cystine-knot motif. An amino acid conservation of 50% or more is marked by a colon. An amino acid conservation of 90% or more is marked by a star.

**Figure S2:**
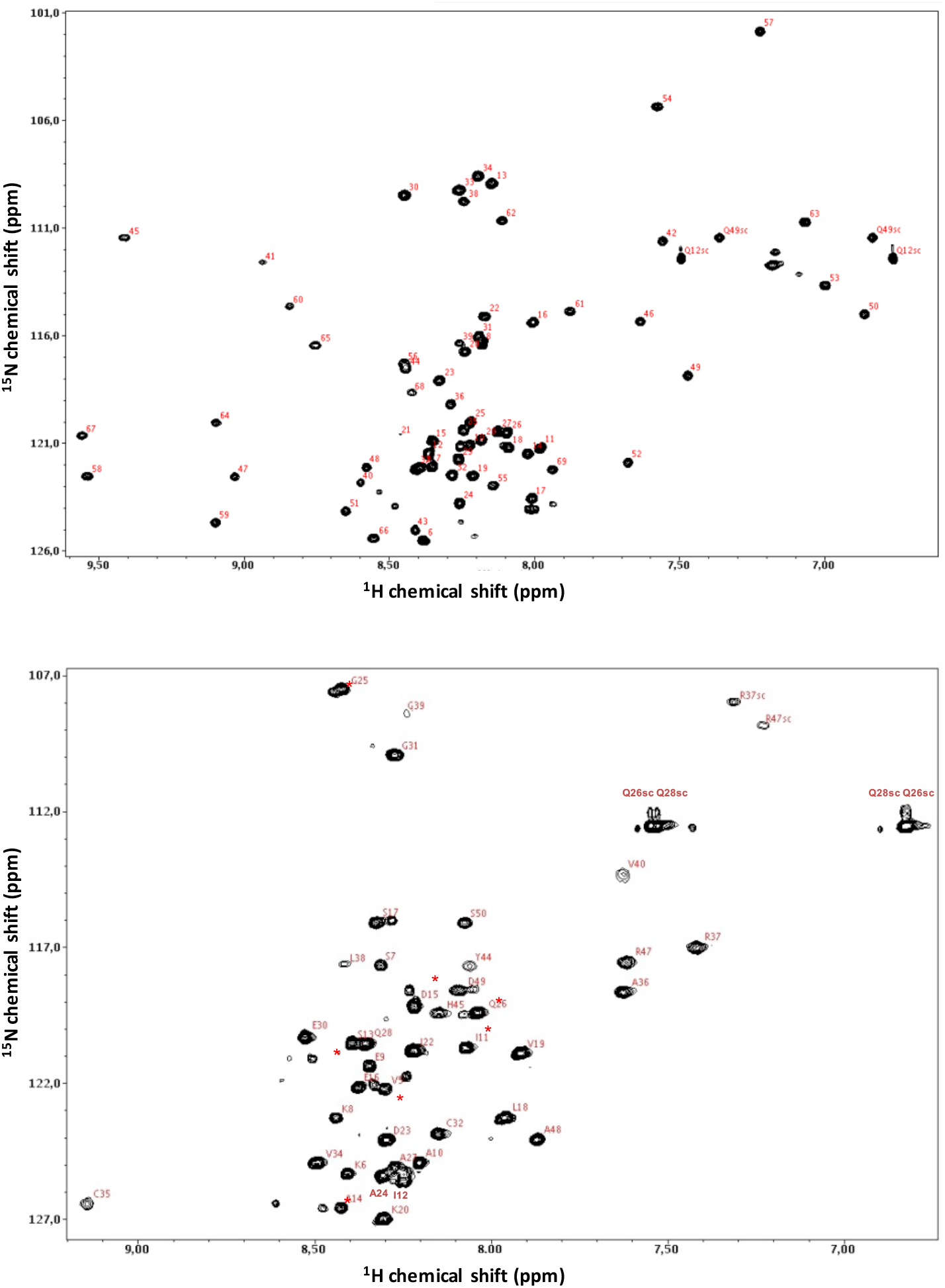
^1^H-^15^N HSQC spectra of recombinant MLP124499. The NMR experiment was recorded on a 600 MHz spectrometer, at 298 K, on a 0.135 mM uniformly ^15^N-labeled sample in phosphate buffer pH 6.0. Backbone amide signals of MLP124499 are labelled with the residue number and « sc » refers to side-chain amide signals. Minor peaks observed in MLP124499 are indicated by asterisks (*).

**Figure S3:**
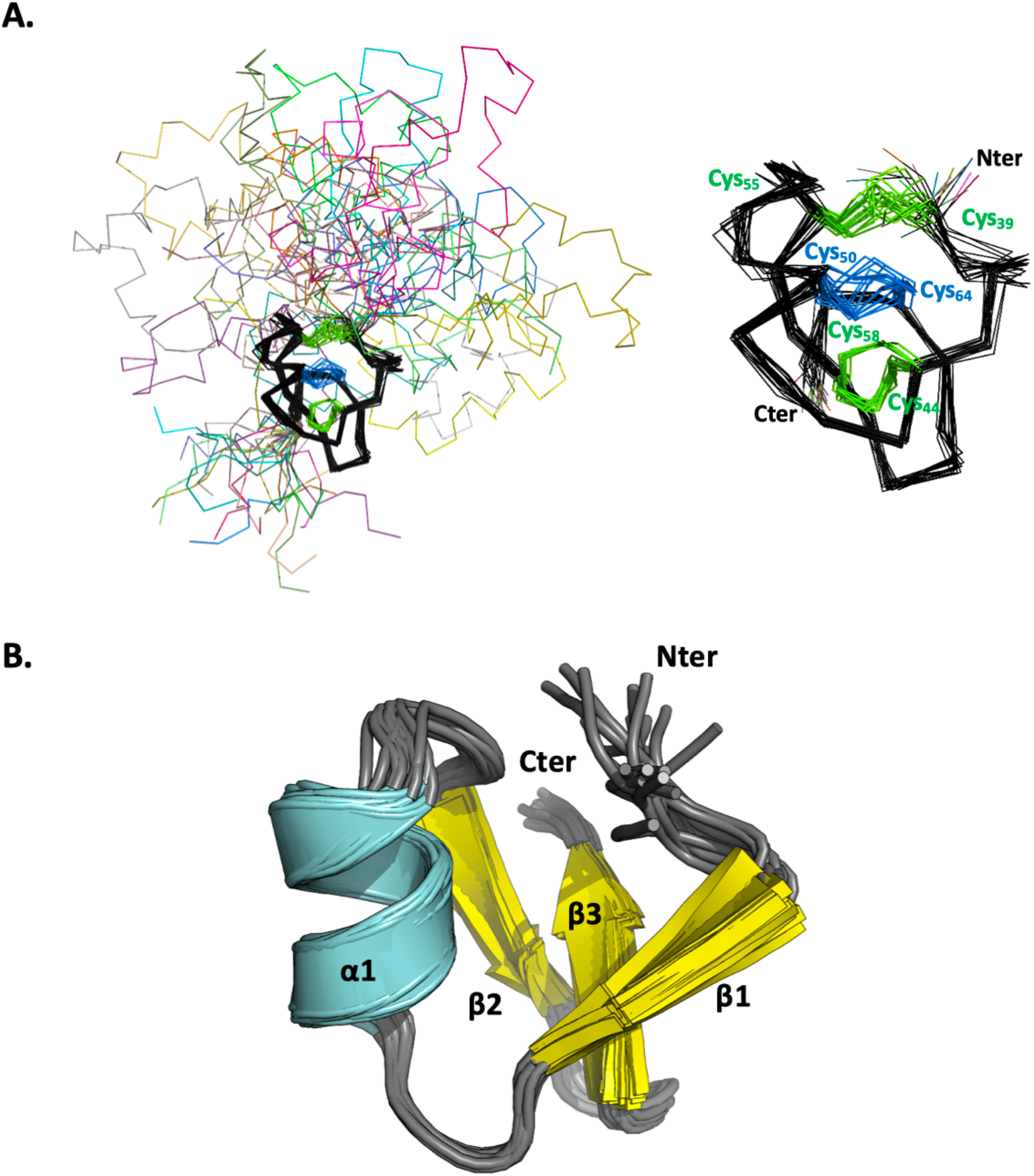
Superposition of the 20 lowest-energy NMR structures of MLP124266. **A.** Superimposition of backbone heavy atoms (N, Cα and C) for a family of 20 lowest energy NMR structures (left) and zoom on cystine-knot (right). Each structure is rendered as ribbon. Cystine-knot (region 39-65) is coloured in black and the three disulfide bridges are labeled and coloured in green (Cys39-Cys55, Cys44-Cys58) or in blue (Cys50-Cys64). Nter and Cter extremities of the cystine-knot are indicated. **B.** NMR models were aligned by their cystine-knot region (region 39-65). Secondary structures are labelled and represented as cartoon. The three disulfide bridges are not represented to clarify the figure. Nter and Cter extremities of the cystine-knot region are indicated.

**Figure S4:**
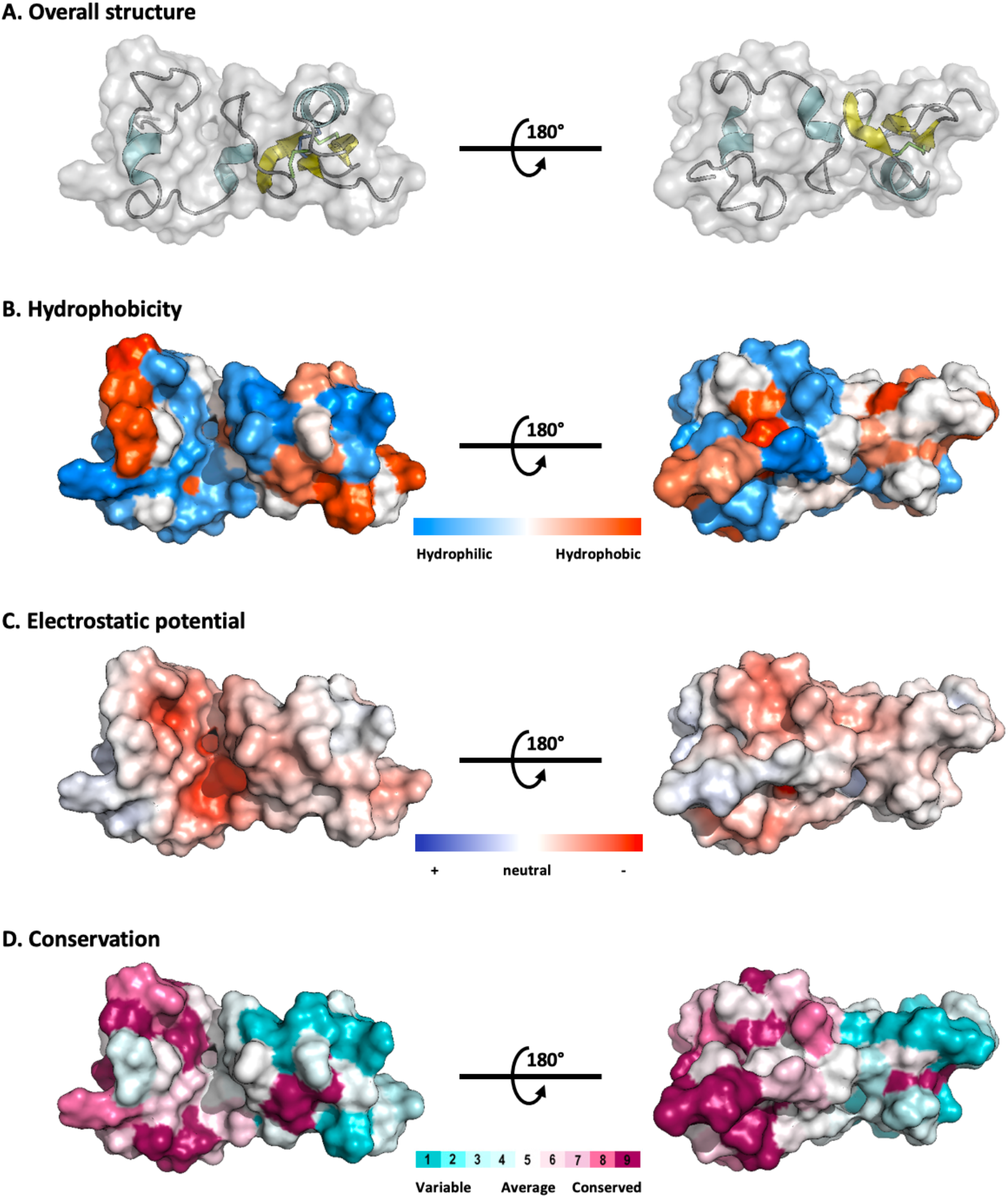
Biophysical properties of MLP124266. Front and rear views on the surface of MLP124266 (**A**) illustrating hydrophobic potential (**B**), the electrostatic Coulomb potential at pH 7.0 using APBS plugin from Pymol 2.0 software with a contour of −10 kT/e to 10 kT/e (**C**) and the conservation of residues generated by Consurf server (**D**). Hydrophobic and hydrophilic patches are shown in orange and in blue, positive-charge and negative-charge densities are coloured in blue and red respectively. Conservation scale ranged from high (purple) to poor conservation (light blue).

**Figure S5:**
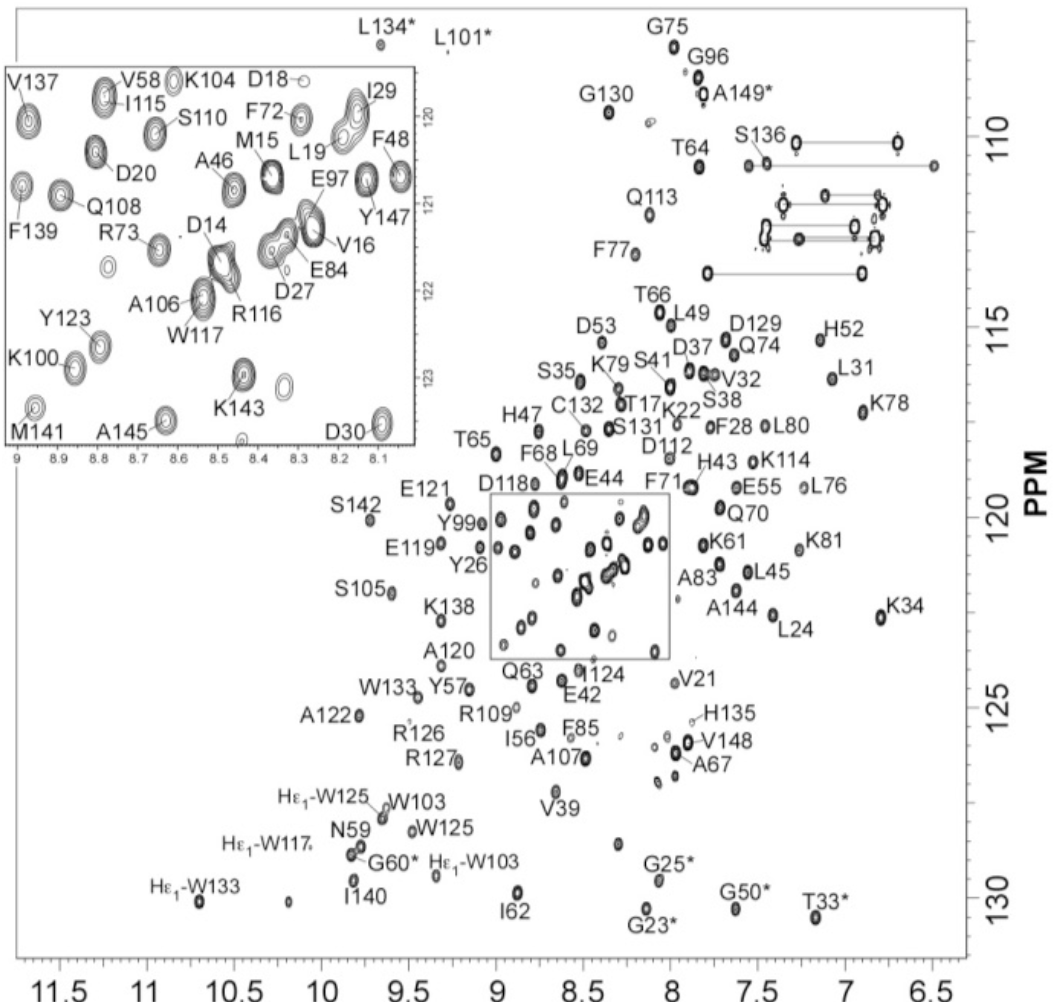
^15^N-HSQC spectra of MLP124017. Cross peak assignments are indicated using the one-letter amino acid and number (the asterisk indicates a folded peak). The central part of the spectrum is expanded in the insert. Missing or non-assigned residues: 1-15, 40, 82, 86-95, 98, 102, 111, 128, 150, 151.

**Figure S6:**
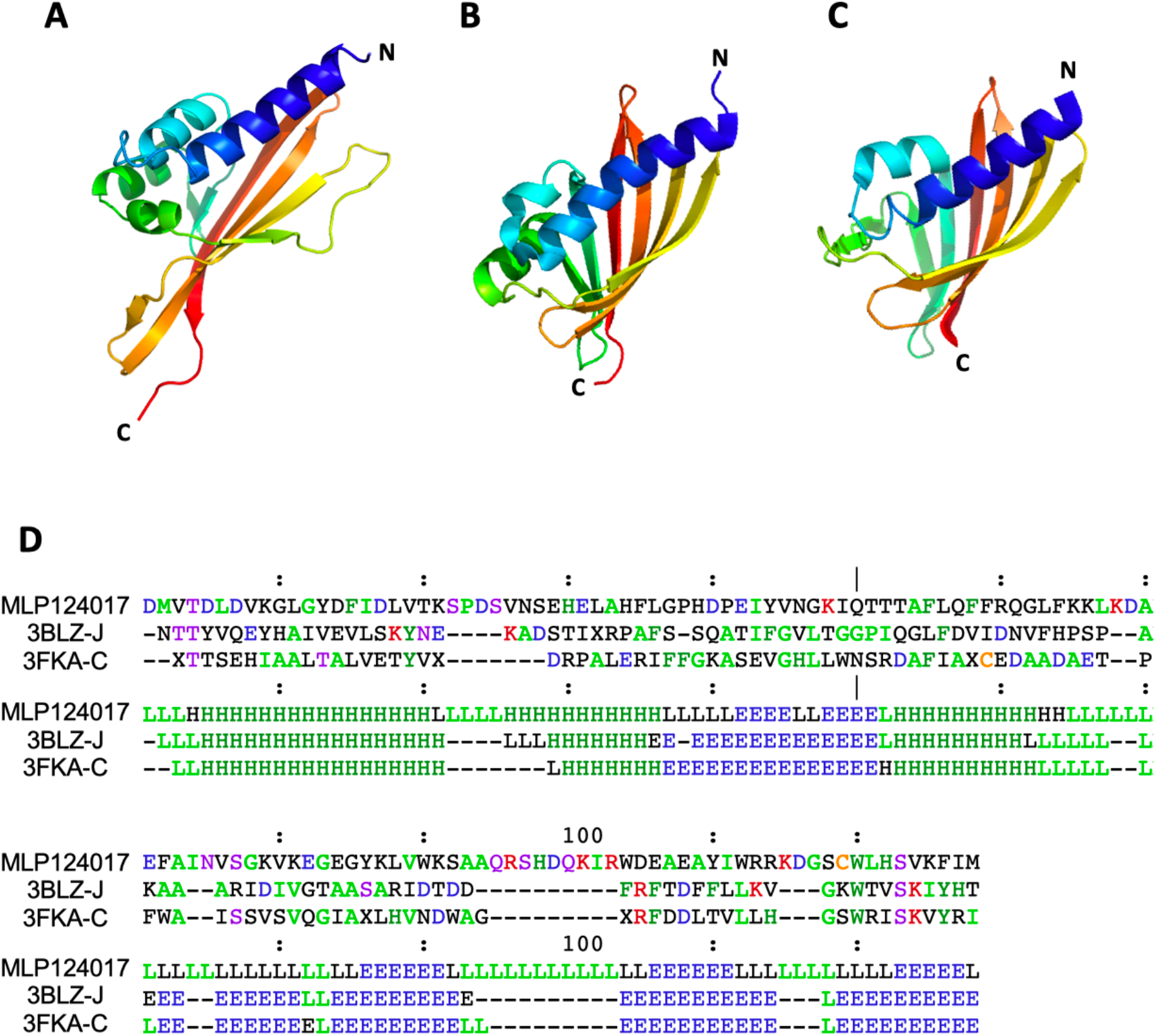
MLP124017 and two bacterial proteins SBAL_0622 and SPO1084 have similar Nuclear-transport factor 2-like fold. Structures of (**A**) MLP124017, (**B**) SBAL_0622 (PDBcode: 3BLZ) and (**C**) SPO1084 (PDBcode: 3FKA) were represented in cartoon with a rainbow coloration using PYMOL (http://www.pymol.org). (**D**) Multiple structural alignment for the three proteins was done with Dali (Holm and Rosenström, 2010) with the MLP124017 as the reference for numbering. The first part shows the amino acid sequences of the selected neighbours. The second part shows the secondary structure assignments by DSSP (H/h: helix, E/e: strand, L/l: coil). The most frequent amino acid type is coloured in each column.

